# Combination of mitomycin C and low-dose metronidazole synergistically against *Clostridioides difficile* infection and recurrence prevention

**DOI:** 10.1101/2024.10.17.617749

**Authors:** Jun-Jia Gong, I-Hsiu Huang, Yuan-Pin Hung, Yi-Wei Chen, Yun-Chien Lin, Jenn-Wei Chen

## Abstract

*Clostridioides difficile* is an anaerobic, spore-forming, Gram-positive pathogen responsible for various conditions from mild diarrhea to severe toxic megacolon and potentially death. Current treatments for *C. difficile* infection (CDI) rely on antibiotics like vancomycin and metronidazole (MTZ); however, the high doses required often disrupt gut microbiota, leading to a high recurrence rate. Mitomycin C (MMC), a chemotherapy drug approved by the FDA, is known for inducing phage production in lysogenic bacterial strains, effectively targeting the host bacteria. Given that 70% of *C. difficile* strains harbor prophages, this study investigates MMC’s potential to enhance antibiotic efficacy against CDI. Our in vitro experiments indicate that MMC acts synergistically with MTZ to inhibit the growth of *C. difficile* strain R20291. Furthermore, this combination decreases biofilm-resident vegetative cell resistance and reduces the MTZ concentration needed to kill *C. difficile* in stool samples ex vivo. In a CDI mouse relapse model, in vivo results show that MMC combined with a low dose of MTZ significantly improves survival rates and reduces fecal spore counts after antibiotic treatment. Overall, these findings suggest that a low-dose combination of MMC and MTZ offers enhanced therapeutic efficacy for CDI, both in vitro and in vivo, and may provide a promising new approach for treatment.

## Introduction

*Clostridioides difficile* is an anaerobic, gram-positive, spore-forming bacterium. The symptoms of *C. difficile* infection (CDI) progress from asymptomatic carriage, diarrhea to severe pseudomembranous colitis, and even death (1). Antibiotics, such as metronidazole (MTZ), vancomycin (VAN), and fidaxomicin, are clinically recommended against CDI (2–4). However, a high dose of antibiotics disrupts the host gut microbiota, leading to recurrent *C. difficile* infection (rCDI) (5). rCDI is a serious clinical challenge, with over 20% of patients experiencing recurrence within 30 days (6), and 60% will relapse again (7). The mortality rate for primary CDI is 2.7%, while for rCDI is 25.4%, which is nearly ten times higher (8). Therefore, investigating effective ways to reduce rCDI caused by antibiotics is crucial.

Mitomycin C (MMC), an FDA-approved anticancer drug, inhibits DNA synthesis by cross-linking DNA (9). In addition to treating several cancer diseases (10–12), it has antibacterial properties (13, 14) and exhibits potent activity against *C. difficile* (15). The combination of two antibiotics is common for treating infections such as tuberculosis and multidrug-resistant bacteria (16).

Since these effects on *C. difficile,* we hypothesize that MMC may assist antibiotic treatment against CDI, and we aim to check the potential role of MMC in CDI treatment. Our study demonstrated that MMC and MTZ synergistically inhibited *C. difficile* RT027 and RT078 strains, reducing the minimal bactericidal concentration (MBC) of MTZ for R20291 biofilm. Additionally, the rCDI animal model trial indicated that combining low-dose MTZ with MMC effectively improved mice survival rates and reduced the spore counts in their stool.

## Materials and Methods

### Bacterial strains and culture conditions

The bacterial strains used in this study are listed in Table S1. All *C. difficile* strains were cultured anaerobically (10% H_2_, 10% CO_2_, and 80% N_2_) at 37℃ in BHI media (Thermo Fisher Scientific, Waltham, MA, USA), alone or with 0.05% L-cysteine (Sigma-Aldrich, St. Louis, USA) and 0.5% yeast extract (Thermo Fisher Scientific). The anaerobic bacteria were maintained in an anaerobic workstation (DG250; Don Whitley Scientific Ltd., West Yorkshire, UK).

### Checkerboard assay

To determine the effects of MMC (Cyrusbioscience, New Taipei City, Taiwan) with MTZ or VAN (Cyrusbioscience), we conducted the checkerboard assay in 96-well plates (SPL life sciences, Pocheon, South Korea) (17). The initial drug concentrations were prepared at 4-fold minimum inhibitory concentration (MIC) values, followed by 2-fold serial dilutions. Next, each well was inoculated with 100 µL of *C. difficile* at 10^6^ colony-forming unit (CFU)/mL in BHIS broth. After 18 hours of incubation at 37°C, the OD_600_ values were measured using the iMark^TM^

Microplate Absorbance Reader (Bio-Rad, Alfred Nobel Drive. Hercules, CA, USA). The combined drug effects were quantified by the fractional inhibitory concentration (FIC) index as follows:

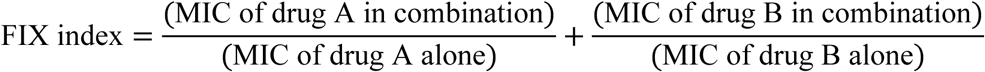

Synergy, FIC index ≤0.5; additive or indifference, FIC index >0.5 but <4; antagonism, FIC index >4 (18). The results represented three independent experiments with the average from triplicate measurements.

### Epsilometer test (E-test) assay

The susceptibility of *C. difficile* to MTZ was determined using the E-test assay (19). The overnight cultures of R20291 were 100-fold diluted using BHIS broth and cultured until the value of OD_600_ reached 1.0. The refreshed cultures were then spread on BHIS agar, with or without MMC, using sterile glass plate spreaders. A single strip of MTZ (Biomérieux, Marcy-l’Étoile, France) was positioned at the center of each plate and incubated at 37°C for 24 hours. The inhibitory concentrations were directly determined at the point of intersection between the inhibition zone and the test strip. This experiment was representative of three independent experiments.

### Biofilm formation and MBC assay

The biofilm formation assay was conducted as previously described (16). In brief, overnight cultures of strain R20291 were 100-fold diluted in BHI broth containing 0.1 M glucose (Avantor, Radnor Township, Pennsylvania, USA) and grown until the OD_600_ value reached 0.1. 200 µL of cultures were inoculated into 96-well plates and anaerobically incubated at 37°C for 72 hours. The supernatants were then replaced with 200 μL of BHI-glucose broth containing antibiotics, followed by a 48-hour incubation. Subsequently, the supernatants were removed, and 200 μL of BHIS broth was used to resuspend the biofilm. The Minimum Bactericidal Concentration (MBC) Assay was performed to determine the efficacy of biofilm disruption (20). Two µL of resuspended biofilm solution were dropped into BHIS agar and incubated at 37°C for 24 hours. This experiment was representative of three independent experiments, each performed in triplicate.

### Ex vivo bacterial-fecal co-culture assay

Fresh feces were collected from C57BL/6 mice and dissociated in sterile ddH_2_O at a ratio of 0.01 g stool per 1 mL ddH_2_O. Overnight cultures of *C. difficile* strain R20291 were refreshed 100-fold in BHIS broth until the OD_600_ reached 1.0. In each group, 5×10^5^ CFU of R20291 in 800 µL BHIS broth were combined, with or without different antibiotics (1 µg/mL of MMC, 0.375 µg/mL of MTZ, 3 µg/mL of MTZ, or 1 µg/mL of MMC+0.375 µg/mL of MTZ). These mixtures were then gently mixed with 200 µL of fecal suspensions and anaerobically incubated for 24 hours. The fecal-bacterial cultures were then diluted 10^6^-fold and spread on selective cycloserine-cefoxitin fructose agar (CCFA) plates for CFU quantification. The experiment was representative of four independent experiments with duplicate measurements. Statistical analysis was performed using One-Way ANOVA with GraphPad Prism 10.0 (GraphPad, La Jolla, CA).

### Spore purification

The spore purification process was performed as previously described (21). Briefly, overnight cultures of *C. difficile* were 100-fold diluted in BHIS broth and grown until the OD_600_ value reached 0.8. The bacterial cultures were then inoculated on SMC agar (22) in 6-well plates (SPL life sciences) and incubated anaerobically at 37℃ for 7 days. *C. difficile* spores were harvested using ice-cold sterile ddH_2_O and stored overnight at 4°C. Spores were centrifuged at 2,330 ×g for 16 min at 4°C and washed five times with ice-cold sterile ddH_2_O to separate spores from cell debris. Spores were purified using Nycodenz (48%; Axis Shield, Oslo, Norway) and centrifuged at 2,500 ×g for 10 min to collect the pellet. After five times wash with ice-cold sterile ddH_2_O, the pellet was resuspended in 1x PBS and stored at 4°C. Purified spores were 10-fold serially diluted and plated on BHIS agar with 0.1% taurocholic acid (Sigma-Aldrich) to determine the CFUs formed by spores.

### Experimental rCDI animal model and sample collection

7-weeks-old male C57BL/6 mice were obtained from the Laboratory Animal Center of NCKU. All mice were managed following the Institutional Animal Care and Use Committee (IACUC) guidelines of NCKU. All animal experiments followed the protocol approved by the IACUC (approval number: NCKU-IACUC-113-068) and the NCKU Biosafety and Radiation Safety Management Division. To determine the effects of combined antibiotic treatment, a rCDI mouse model was performed as previously described (23). Briefly, mice were intraperitoneally injected with 10 mg/kg of clindamycin 24 hours before infection, then orally challenged with 100 µL containing 10^4^ CFU R20291 spores. Different antibiotic treatments were orally given on days 2 and 3 post-infection. To determine whether MMC can synergically assist low-dose of MTZ against *C. difficile* infection and recurrence, mice were divided into six groups and respectively treated with (1) sterile 1x PBS (Mock group, 3 mice and R20291 group, 8 mice), (2) 50 mg/kg of MTZ dissolved in 1x PBS (MTZ_HC_ group, 8 mice), (3) 7 mg/kg of MTZ dissolved in 1x PBS (MTZ_LC_ group, 8 mice), (4) 5 mg/kg of MMC dissolved in 1x PBS (MMC group, 8 mice), and (5) 7 mg/kg of MTZ combined with 5 mg/kg MMC dissolved in 1x PBS (MTZ_LC_+MMC group, 8 mice). In the animal experiment to compare the efficacy of MTZ_LC_+MMC treatment and VAN treatment, mice were divided into three groups and respectively treated with (1) 50 mg/kg of MTZ dissolved in 1x PBS (MTZ_HC_ group, 9 mice), (2) 20 mg/kg of VAN dissolved in 1x PBS (VAN group, 9 mice), (3) 7 mg/kg of MTZ combined with 5 mg/kg MMC dissolved in 1x PBS (MTZ_LC_+MMC group, 9 mice). During the infection, mice were monitored daily for clinical sickness score (CSS) (24), and fecal samples were collected to quantify the number of spores on days 3, 5, and 7 post-infection. Mice sera were analyzed for glutamic pyruvic transaminase (GPT), glutamic oxaloacetic transaminase (GOT), blood urea nitrogen (BUN), and creatinine (CRE) using NX-600 dry biochemistry analyzer (Fujifilm, Minato, Tokyo, Japan). After colonic morphology was observed, harvested organs were fixed in 10% formaldehyde (BS Chemical, Kretinga, Lithuania) or stored at −80°C for further analysis. Statistical analysis was performed using GraphPad Prism 10.0, and significance was determined by t-test and One-Way ANOVA.

### Quantification of *C. difficile* spore-forming CUFs

Fresh fecal samples collected on days 3, 5, and 7 post-infection were weighted and diluted by 1 mL of ddH_2_O with 100 mg of 0.1 mm zirconia beads (BioSpec Products, Inc., Bartlesville, Oklahoma, USA). After the homogenization using the BeadBeater (Bertin Technologies, avenue Ampère, Montigny-le-Bretonneux, France) for 50 sec (23), fecal samples were heat-killed at 65°C for 20 min and plated on BHIS agar containing 0.1% TCA to quantify the CFUs. The plates were anaerobically incubated at 37°C for 24 hours and CFU numbers were calculated after normalizing to the fecal weight. This experiment was representative of three independent experiments and each in duplicate. Statistical analysis was performed using GraphPad Prism 10.0, and significance was determined by t-test and One-Way ANOVA.

## Results

### Mitomycin C showed a synergistic antibacterial activity in combination with metronidazole against *C. difficile* RT027 strain R20291 and RT078 clinical strains

The previous study used MMC to induce phage production in *C. difficile* and promote the lytic cycle (21). Interestingly, over 70% of *C. difficile* strains naturally carry prophages (25). Therefore, we wanted to check whether MMC could be combined with VAN or MTZ as a novel CDI therapeutic strategy. First, we performed the checkerboard assay (Figure 1A), and the MIC of MMC and MTZ were determined to be 2 µg/mL and 3 µg/mL, respectively (Figure S1A). The combination of MTZ with MMC showed synergistic antibacterial activity in inhibiting the growth of R20291, as the FIC index was less than 0.5 (Figure 1B). MTZ combined with 1 µg/mL MMC could reduce the MIC of MTZ for inhibiting *C. difficile* growth from 3 µg/mL to 0.375 µg/mL (an 8-fold reduced MIC value) (Figure S1A). While the combination of VAN and MMC exhibited an additive activity only (Figure 1B). Furthermore, the MTZ-MMC treatment also synergistically against RT027 and RT078 *C. difficile* clinical strains (Figure 1C); MMC additively assisted VAN in inhibiting RT078 strains (Figure S1B). These results demonstrated that MMC enhances the effects of CDI-treating antibiotics.

**Figure 1.**
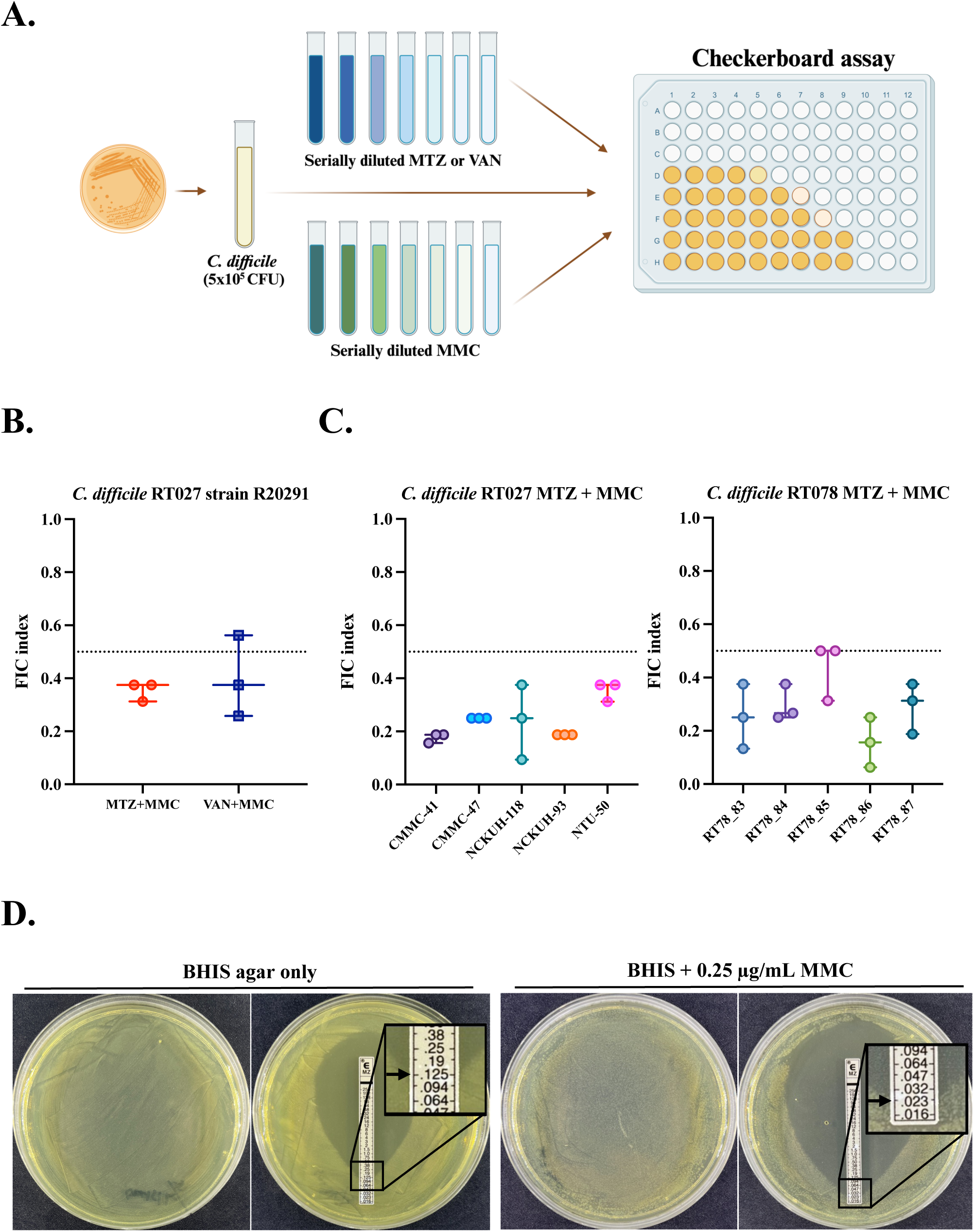
MMC and MTZ demonstrated synergistic inhibition against specific *C. difficile* strains, including RT027 (e.g., R20291) and RT078 in vitro. (**A**) Flowchart of checkerboard assay (Created with BioRender.com). FIC index of MMC combined with MTZ or VAN co-culturing with (**B**) *C. difficile* strain R20291, (**C**) RT027 and RT078 clinical strains. The MIC of MTZ alone or a combination of MMC with MTZ against *C. difficile* was determined using E-test strips (**D**). MTZ, metronidazole. VAN, vancomycin. MMC, mitomycin C. FIC index, fractional inhibitory concentration index. MIC, minimum inhibitory concentration. The results represented three independent experiments with the average from triplicate measurements.

E-test strips were used to confirm the synergistic effects of MTZ and MMC for combating R20291. The MIC of MTZ strip alone was 0.125 µg/mL (Figure 1D, left). Notably, 0.25 µg/mL of MMC would not influence the growth of bacteria (Figure 1D, middle), but effectively reduced the MIC of MTZ to 0.023 µg/mL (a 5.4-fold reduced MIC value; Figure 1D, right). These in vitro experimental results indicate that the combination of MTZ and MMC can more effectively inhibit *C. difficile* strain R20291 than MTZ alone.

### Mitomycin C could effectively enhance the metronidazole effect on the strain R20291 biofilms

In the CDI mouse model, *C. difficile* strain R20291 can aggregate and form a glycan-rich biofilm, which is associated with increased antibiotic resistance and rCDI (26–28). Therefore, we tried to clarify whether MMC can help MTZ reduce the required concentration against biofilm by the MBC assay (Figure 2A). As shown in Figure 2B, the MBC of MTZ for R20291 vegetative cells was 3 µg/mL. However, the MBC of MTZ required to eliminate R20291 in biofilm increased to 18 µg/mL (Figure 2C, left). Interestingly, combining MTZ with 1 µg/mL of MMC markedly decreased the MBC of MTZ to 9 µg/mL (Figure 2C, middle). MMC showed no toxicity on R20291 vegetative cells until the concentration reached 4 µg/mL (Figure 2C, right). To determine whether any vegetative cells were released outside the biofilm, we dropped the biofilm supernatant on BHIS agar plates. As shown in Figure S2, no lived bacterium was detected outside the biofilm after treatment with antibiotics. Taken together, MMC could enhance MTZ to effectively reduce its MBC against R20291 biofilm.

**Fig 2.**
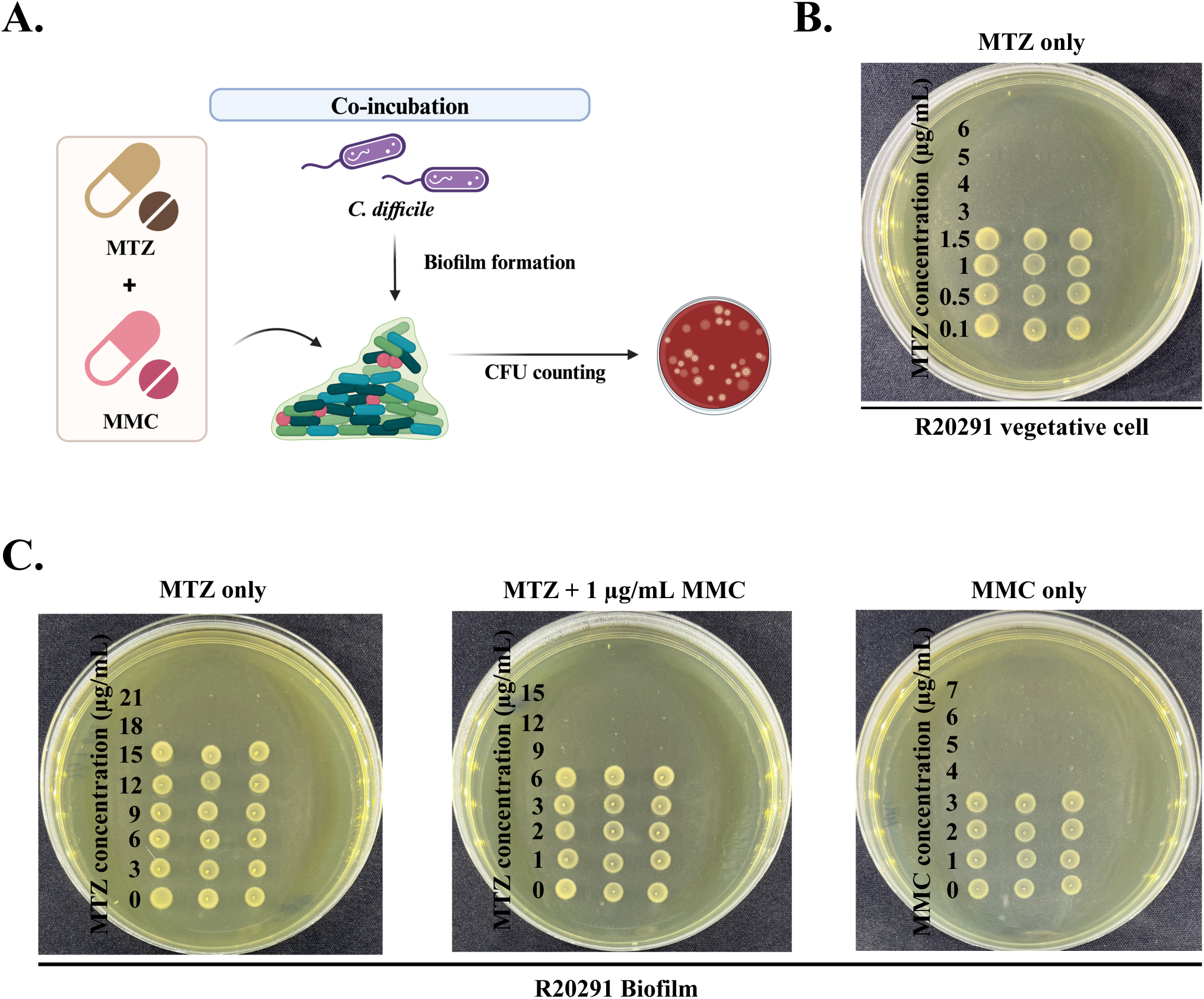
Combining MMC and MTZ therapy effectively reduced the antibiotic resistance of R20291 biofilms. (**A**) The flowchart of antibiotic treatments against *C. difficile* biofilm formation (Created with BioRender.com). (**B**) MBC assay was used to determine the concentration of MTZ that can kill R20291 vegetative cells. After 3 days of incubation to form biofilms, the MBC concentration of (**C**) MTZ alone, MTZ combined with 1 µg/mL MMC, and MMC alone to combat R20291 within the biofilm was determined. MBC, minimum bactericidal concentration. This experiment was representative of three independent experiments, each performed in triplicate.

### Combination of mitomycin C with metronidazole inhibits *C. difficile* colonization in fresh mouse feces

To simulate the diverse microbial gut environment, *C. difficile* strain R20291 was mixed with fresh mouse fecal suspensions to assess the efficacy of a low concentration of MTZ combined with MMC against *C. difficile* ex vivo. After co-culturing for 24 hours, 10^6^ dilutions of bacterial-fecal cultures were spread on BHIS agar to assess the bacterial CFU count (Figure 3A). As shown in Figure 3B, no CFU was observed in the 3 µg/mL of MTZ-treated group, indicating that the number of *C. difficile* was less than 10^6^ CFU/mL. With the combination treatment of 0.375 µg/mL MTZ and 1 µg/mL MMC, an average of 4.5×10^6^ CFU/mL was observed. By comparison, the average CFU counts with 0.375 µg/mL of MTZ (MTZ_LC_) or 1µg/mL of MMC alone were 7.8×10^7^ CFU/mL and 4.9×10^7^ CFU/mL, respectively. The difference may be due to the low concentration of antibiotic treatments, which affect other bacteria competing with *C. difficile* while having no direct impact on *C. difficile* in the mouse feces. In summary, these results demonstrated that combining MTZ with MMC is a potential strategy to inhibit *C. difficile* colonization ex vivo.

**Fig 3.**
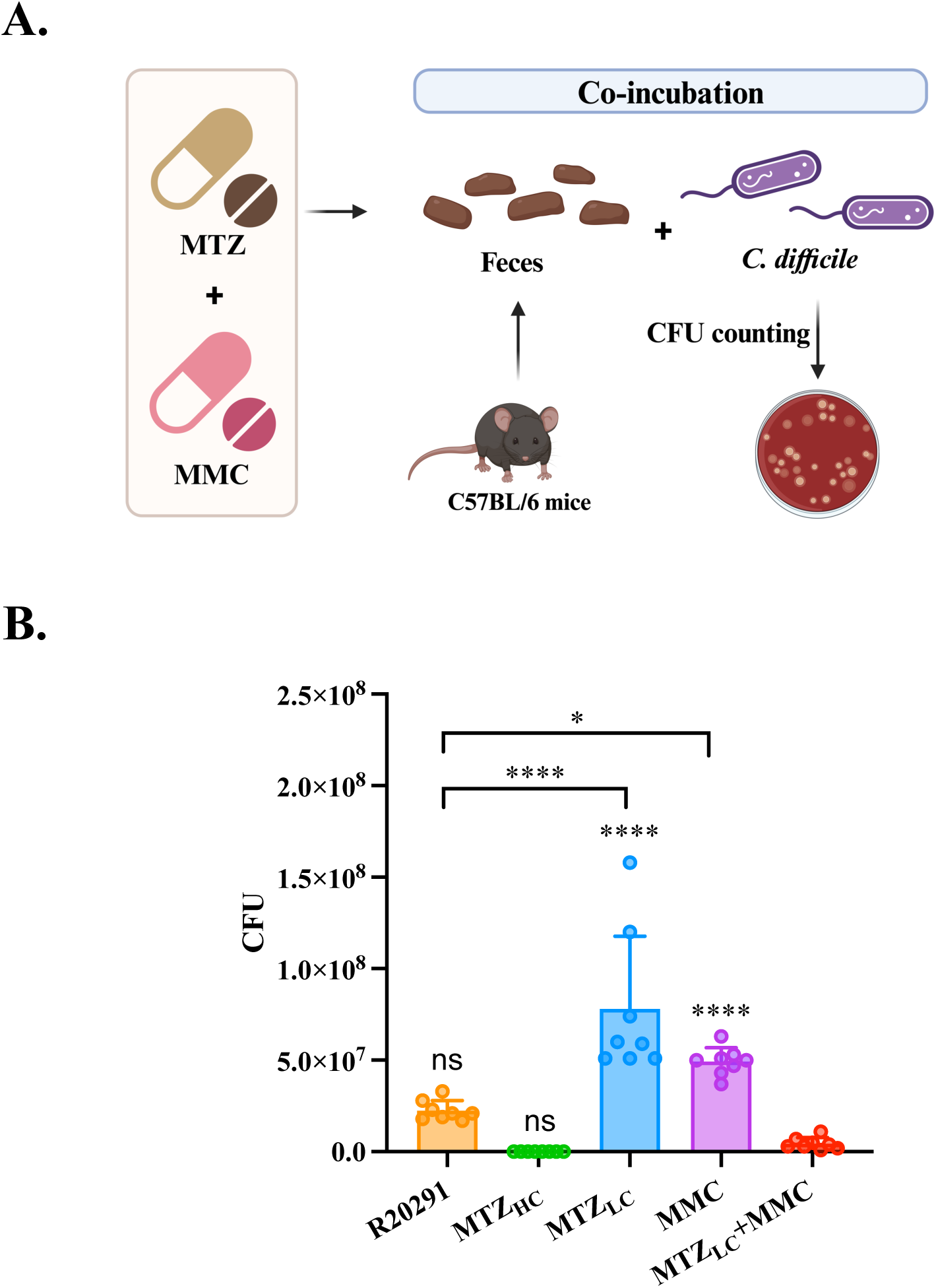
The combination of MMC and low-dose MTZ significantly inhibited the growth of R20291 in mouse feces. (**A**) Ex vivo model of R20291 co-cultured with mouse feces and treated with different kinds of antibiotic treatment (Created with BioRender.com). (**B**) Fecal-bacterial mixtures cultured for 24 hours were 10^6^-fold diluted, and the CFU counts of *C. difficile* were determined by spreading on CCFA plates. CFU, colony-forming units. MTZ_HC_, high concentration of MTZ (3 µg/mL). MTZ_LC_, low concentration of MTZ (0.375 µg/mL). MMC, 1 µg/mL MMC. MTZ_LC_+MMC, 0.375 µg/mL MTZ combined with 1 µg/mL MMC. The experiment was representative of four independent experiments with duplicate measurements. Statistical analysis was performed using the one-way analysis of variance test (\**p* ≤ 0.05; \*\*\*\**p* ≤ 0.0001; ns, not significant)

### Treating metronidazole with mitomycin C successfully increased the survival rate and decreased the spore number in a recurrence CDI mouse model

Given that the combination of low-dose MTZ and MMC effectively inhibited *C. difficile* in vitro and ex vivo, we want to determine whether this strategy can be applied in vivo (Figure 4A). We referred to the rCDI mouse model (23), and challenged mice with purified R20291 spores (Figure S3A, B). Only the non-infected and infected group treated with MTZ_LC_+MMC demonstrated a 100% survival rate, while other groups consistently fell below 80% (Figure 4B, left). Furthermore, both R20291 group and MMC group exhibited the lowest weight on day 2 post-infection. However, by day 7 post-infection, all surviving mice had recovered to over 95% of their initial weight (Figure 4B, right).

**Fig 4.**
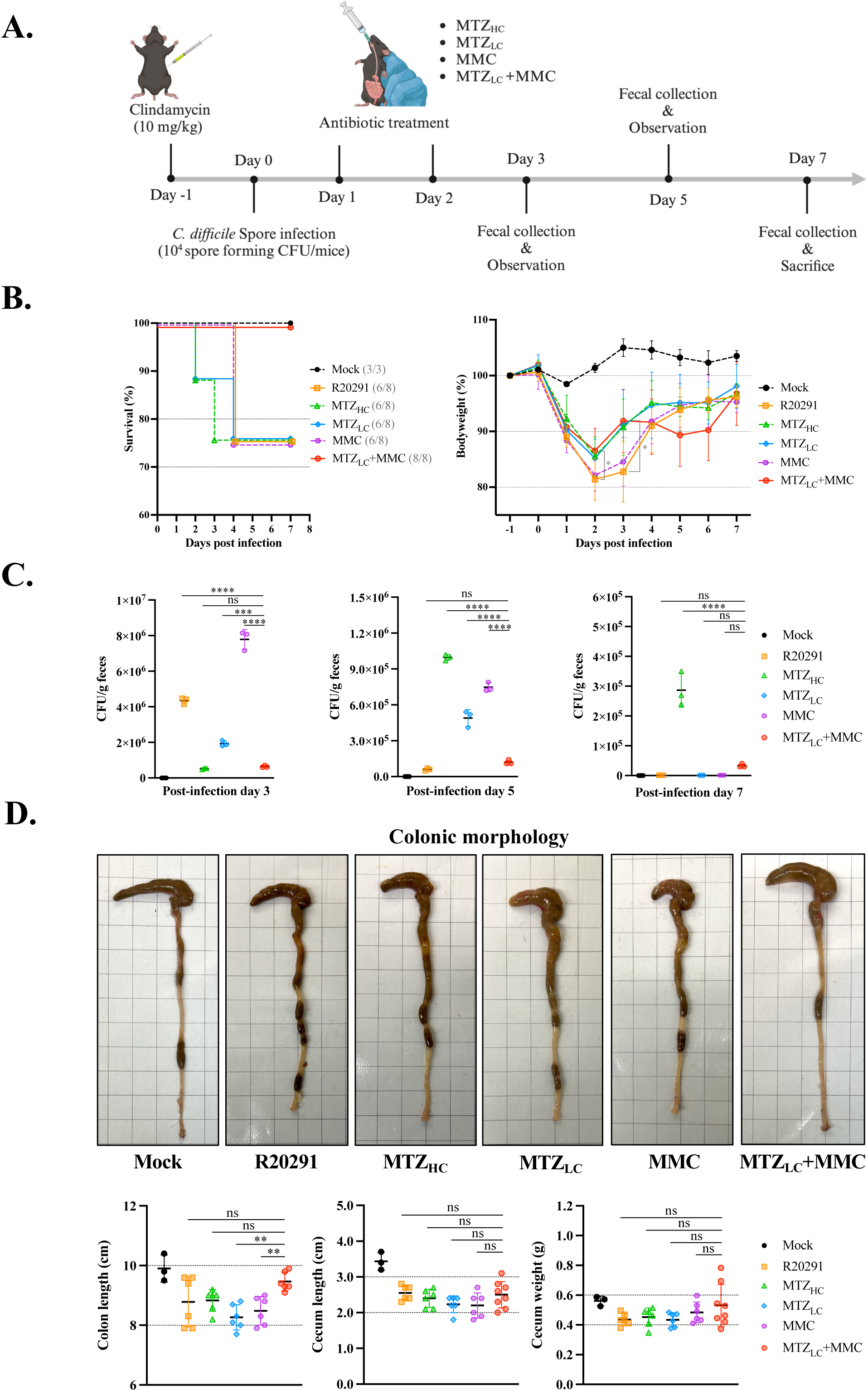
Application of combining MMC with low-dose MTZ treatment in the mouse CDI recurrence model. (**A**) Schematic diagram of the mouse CDI recurrence model (Created with BioRender.com). C57BL/6 mice were given clindamycin (10 mg/kg) and orally challenged with 10^4^ CFU spores. After two days of treatment with (n=8/group) or without (Mock, n=3; R20291, n=8) antibiotics, the severity of infection symptoms was observed, and feces were collected to assess the number of spores. (**B**) The survival and body weight change rate of mice were monitored daily from the day of infection (pi-day 0) until being sacrificed. (**C**) The CFU of *C. difficile* spore was determined using mouse feces collected on the days 3, 5, and 7 post-infection. (**D**) The morphology and quantification of cecal and colonic tissues. The colon length, cecum length, and cecum weight were harvested when the mice were sacrificed on day 7 post-infection. Each point represents the data of one mouse. Statistical analysis was performed using GraphPad Prism 10.0, and significance was determined by the one-way analysis of variance (**≤ 0.01; ***≤ 0.001; \*\*\*\**p* ≤ 0.0001; ns, not significant)

To determine the recurrence of *C. difficile* in mice, we collected feces on days 3, 5, and 7 post-infection and plated them on BHIS agar with 0.1% taurocholic acid (TCA). The spore-formed CFU counts showed no significant difference between MTZ_HC_ and MTZ_LC_+MMC groups on day 3 post-infection (Figure 4C, left). However, the CFU counts in other infected groups were significantly higher than those in the MTZ_LC_+MMC group (Figure 4C). On day 5 post-infection, the spore-formed CFU counts in R20291 and MTZ_LC_+MMC groups were significantly lower than in other infected groups (Figure 4C, middle). Notably, the MTZ_HC_ group had a significantly highest CFU number among all groups (Figure 4C, middle). On day 7 post-infection, despite a decrease in CFU number from all infected groups, the CFU number in the MTZ_HC_ group remained significantly higher than that in the MTZ_LC_+MMC group (Figure 4C, right). During the CDI, the daily clinical sickness score of mice in the MTZ_LC_+MMC group was lower than other infection groups (Figure S3C). The colonic morphology of infected mice was an important indicator for assessing intestinal inflammation (29). As shown in Figure 4D, all surviving mice recovered at day 7 post-infection with no significant difference in their cecum length and weight. The colon length of the MTZ_LC_+MMC group was significantly higher than that of the MTZ_LC_ and MMC groups. Taken together, the rCDI in vivo experiments demonstrated that the MTZ_LC_+MMC treatment could express the highest mouse survival rate and prevent the relapse of CDI.

### Synergistic effects of mitomycin C and metronidazole enhanced mice survival rate and reduced stool spore number compared to vancomycin treatment

Previous studies suggest that VAN has superior cure rates for severe CDI and better outcomes for infected mice than those treated with MTZ (30, 31). However, VAN treatment was associated with a significant disruption of the intestinal microbiota, thereby increasing the risk of the rCDI (32, 33). Therefore, we wanted to assess whether combining MTZ with MMC could express a better effect than VAN in the rCDI mouse model.

After being infected with R20291 spores, mice received VAN, MTZ_HC_, or MTZ_LC_+MMC treatments, respectively (Figure 5A). The MTZ_LC_+MMC treatment provided a higher survival rate (89%) compared to the VAN (67%) or MTZ_HC_ (67%) groups (Figure 5B, left). In body weight change rate, VAN and MTZ_HC_ groups exhibited an appreciable decrease on day 3 to 4 post-infection; this trend was less obvious in MTZ_LC_+MMC group (Figure 5B, right), all groups of mice recovered to over 90% of their initial weight on day 7 post-infection (Figure 5B, right). Although the CFU counts of spore in feces from MTZ_LC_+MMC group were significantly higher than the other groups on day 3 post-infection (Figure 5C, left). However, on days 5 and 7 post-infection, the CFU counts in MTZ_LC_+MMC group were significantly lower than other two groups (Figure 5C, middle and right). In addition, the VAN group had increased nearly ten times spore number in their feces on day 5 post-infection (Figure 5C, middle), while the MTZ_HC_ group maintained a high spore level in feces from day 3 to 7 post-infection (Figure 5C). In mice intestinal inflammation, except for the cecal weight in MTZ_LC_+MMC group was significantly higher than in the MTZ_HC_ group; mice cecum and colon morphology showed no significant difference between each group (Figure 5D). Finally, to ensure that combining low-dose MTZ with MMC for CDI would not cause liver and kidney damage in mice. Serum was collected on day 7 post-infection and analyzed for glutamic pyruvic transaminase (GPT), glutamic oxaloacetic transaminase (GOT), blood urea nitrogen (BUN), and creatinine (CRE) using a dry biochemical analyzer, and the values of each indicator exhibited no significant difference between the groups (Figure 5E). In summary, our results demonstrated that combining low-dose MTZ with MMC showed the highest survival rate and effectively reduced intestinal spore count in CDI mice compared to VAN or MTZ-treated groups.

**Fig 5.**
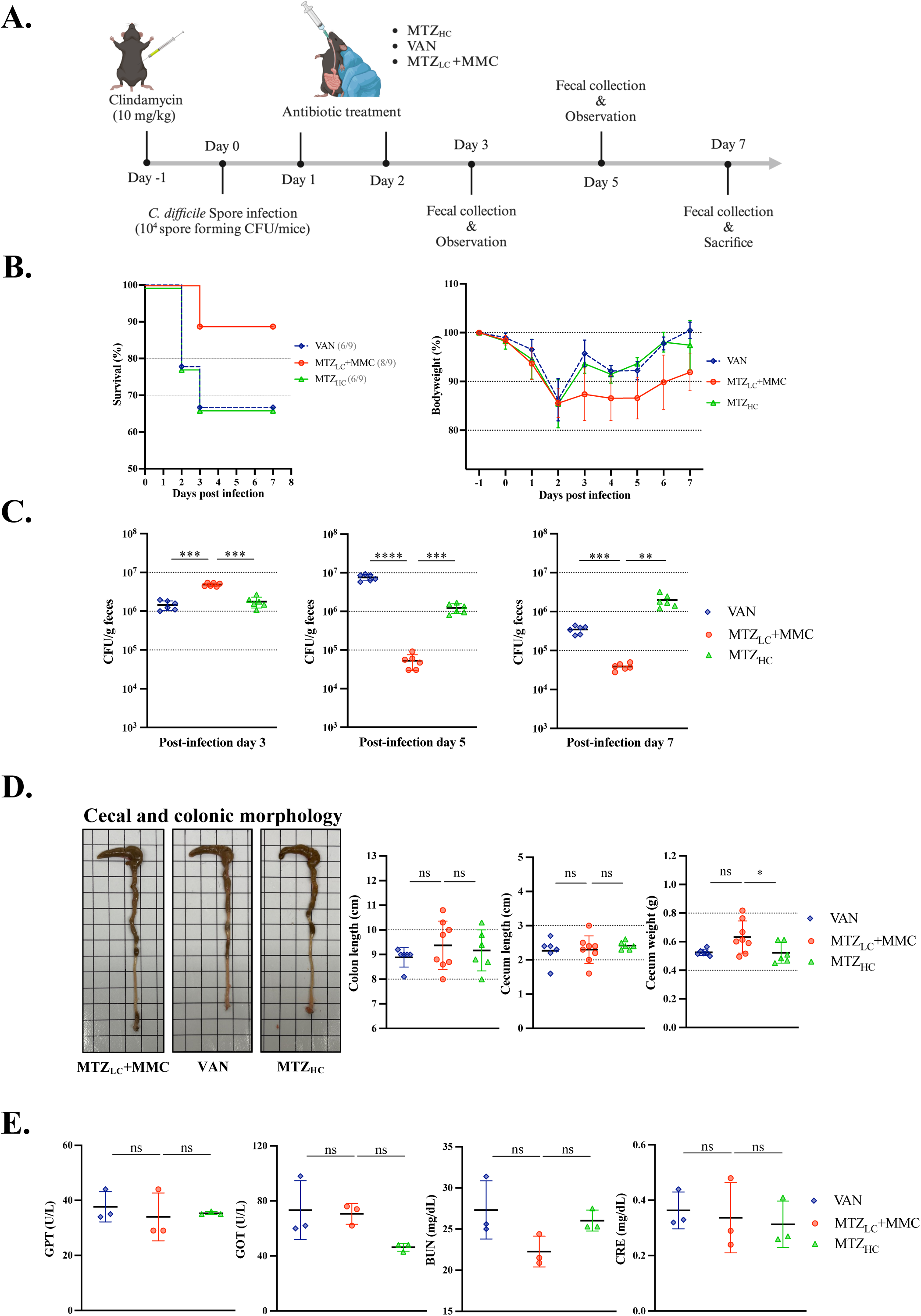
Comparing the effect of MMC-MTZ_LC_ treatment and clinical antibiotics against *C. difficile* infection and recurrence in vivo. (**A**) Diagram of the mouse CDI recurrence model (Created with BioRender.com). Mice (n = 9/group) were orally challenged with *C. difficile* spores, followed by two days of antibiotic treatment. (**B**) Mice survival rate and the daily percentage of mouse body weight were recorded after being infected with R20291 spores. (**C**) Mice feces were collected on day 3, 5, and 7 post-infection. The homogenized feces were diluted and plated on CCFA plates to measure the spore-forming CFU counts. (**D**) The morphology of mice cecum and colon. The degree of intestinal inflammation was evaluated by quantifying the colonic length, cecum length, and weight. Each point represents the data of a mouse. (**E**) The values of GPT, GOT, BUN, and CRE in mice serum. The dry biochemical analyzer was used to measure various indicators in the serum to evaluate the damage and impact of different treatment processes on the liver and kidney functions of mice. GPT, glutamic pyruvic transaminase. GOT, glutamic oxaloacetic transaminase. BUN, blood urea nitrogen. CRE, creatinine. Statistical analysis was performed using GraphPad Prism 10.0, and significance was determined by the t-test and one-way analysis of variance (*≤ 0.05; **≤ 0.01; ***≤ 0.001; \*\*\*\**p* ≤ 0.0001; ns, not significant)

## Discussion

The CDI patients still rely on VAN and MTZ treatments (2–4), however, these antibiotics disrupt gut microbiota to cause the recurrence (34). In this study, we found that MMC can synergistically cooperate with MTZ against *C. difficile*. Moreover, MMC can boost MTZ’s effectiveness in eliminating *C. difficile* within biofilms, despite biofilm formation providing protection for *C. difficile* against antibiotic attacks, which can result in recurrent infections (26–28, 35). Fidaxomicin can penetrate biofilms and disrupt their structure, thereby enhancing the bactericidal activity (36, 37). Future work should observe the biofilm morphology to determine whether the MTZ-MMC treatment directly disrupts biofilms to kill *C. difficile* vegetative cells. Our in vivo experiments showed that combining MTZ_LC_ with MMC increased mice survival rates after the spore challenge. However, the recovery of body weight in this group was not as good as in other infected groups. Most mice of other groups with severe body weight loss have died in CDI, leading to the lived mice having a higher average body weight than the MTZ_LC_+MMC group. The CFUs formed by spore in mice fecal samples to evaluate CDI recurrence showed no significant difference between the R20291 and MTZ_LC_+MMC groups after day 5 post-infection. Colonization resistance is critical for recovery and preventing rCDI through the healthy microbiota in inhibiting *C. difficile* colonization and overgrowth (38–40). However, antibiotics disrupt microbiota diversity (41). Accordingly, we consider that mice in both groups might suffer less damage in their microbiota and recover more rapidly to prevent *C. difficile* colonization. Nevertheless, further analysis of intestinal microbial composition between each group is needed to confirm our hypothesis.

MTZ disrupts DNA synthesis (42), while MMC cross-links DNA to prevent bacterial replication (43). Our study indicated that combining MTZ with MMC provides better action in inhibiting bacterial DNA replication against *C. difficile* infection.

In summary, our findings reveal a novel therapeutic strategy that MMC enhances the efficacy of MTZ against CDI. The overexpression of NimB in some mutant strains reduces the ability of MTZ to treat CDI, leading to antibiotic resistance (44). This issue did not occur in our MTZ-MMC combination therapy for *C. difficile*. Furthermore, the low dose of MTZ combined with MMC may effectively decrease rCDI by minimizing gut microbiota disruption (38, 40). Nevertheless, further experiments are necessary to confirm the safety, optimal dosage, and potential side effects of the combined antibiotic therapy before the clinical application.

## Acknowledgments

This work was supported by the National Science and Technology Council (Grants: 113-2314-B-006-043-MY3; awarded to J. W. C.). The funding agency played no role in study design, data collection or analysis, manuscript preparation, or decision to publish. We thank the Laboratory Animal Center, College of Medicine at National Cheng Kung University (NCKU), Taiwan, accredited by AAALAC International and Taiwan Animal Consortium. J. J. G. contributed to the methodology, validation, formal analysis, investigation, visualization, and original manuscript writing. I. H. H. contributed to conceptualization and methodology. Y. P. H. contributed to conceptualization, methodology, and resources. Y. W. C. contributed to conceptualization, methodology, and visualization. Y. C. L. contributed to investigation. J. W. C. contributed to the conceptualization, methodology, supervision, project administration, funding acquisition, and manuscript editing. All authors declare that they have no conflicts of interest.

