## Supplemental data for "Combination of mitomycin C and low-dose metronidazole synergistically against *Clostridioides difficile* infection and recurrence prevention"

**Supplementary Information**

**Figure S1. In vitro evaluation of MMC assisted antibiotics against *C. difficile* growth.** (A) The checkerboard assay was performed to determine bacterial growth after being treated with different antibiotic concentrations. (B) VAN+MMC efficacy inhibited RT078 clinical strains was evaluated using the checkerboard assay.

**Figure S2. No viable bacteria detected in extracellular biofilm after treatment with MTZ and MMC.**

**Figure S3. Preparation of CDI recurrence model and observation of mouse health status.** (A) Nycodenz-purified R20291 spores were stained with malachite green. Green: R20291 spores. Pink: R20291 vegetative cell debris. (B) Spores were germinated on 0.1% taurocholic acid BHIS agar to calculate the CFU counts. (C) Daily monitoring of mouse clinical sickness score (CSS) from the 3 to 7 days post infection.

### Supplemental figures

#### Fig S1A.

| MTZ (µg/mL) | | | | | | | | | | | | | |
|---|---|---|---|---|---|---|---|---|---|---|---|---|---|
| 6 | 11 | 10 | 10 | 11 | 11 | 11 | 11 | 11 | 11 | 11 | 11 | 10 | |
| 3 | 12 | 12 | 11 | 11 | 11 | 11 | 11 | 11 | 11 | 11 | 11 | 10 | |
| 1.5 | 64 | 70 | 65 | 67 | 54 | 61 | 67 | 12 | 12 | 12 | 13 | 14 | |
| 0.75 | 91 | 99 | 98 | 97 | 97 | 97 | 87 | 68 | 11 | 12 | 13 | 14 | |
| 0.375 | 91 | 91 | 94 | 93 | 91 | 98 | 77 | 53 | 11 | 12 | 13 | 15 | |
| 0.1875 | 111 | 101 | 103 | 100 | 100 | 100 | 87 | 86 | 24 | 12 | 13 | 15 | |
| 0.09375 | 102 | 108 | 107 | 105 | 105 | 102 | 95 | 98 | 83 | 12 | 14 | 15 | |
| 0.046875 | Drug Free | 103 | 116 | 105 | 104 | 102 | 91 | 91 | 95 | 11 | 12 | 14 | |
| | 0.004 | 0.078 | 0.016 | 0.031 | 0.063 | 0.125 | 0.25 | 0.5 | 1 | 2 | 4 | 8 | MMC (µg/mL) |

#### Fig S1B.

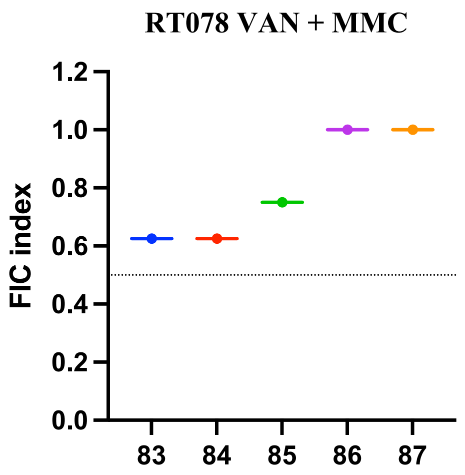


#### Fig S2.

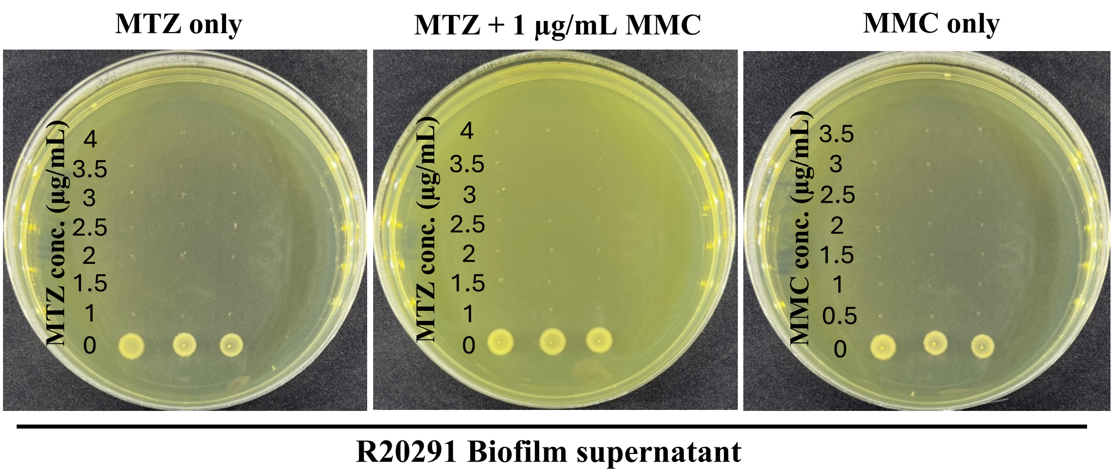

#### Fig S3A.

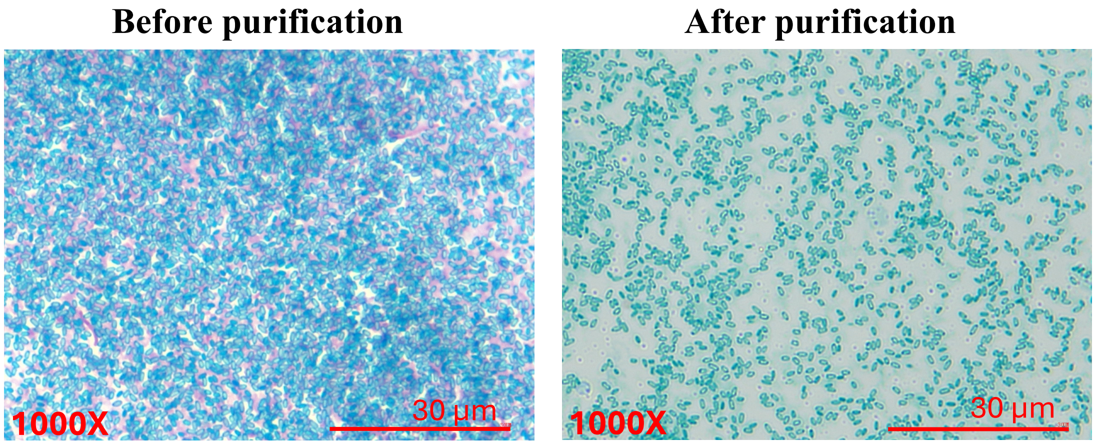


#### Fig S3B.

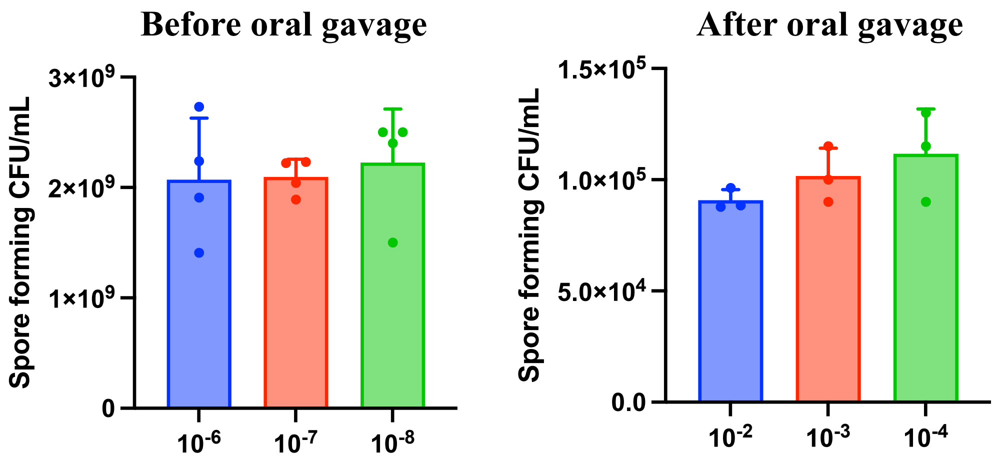


#### Fig S3C.

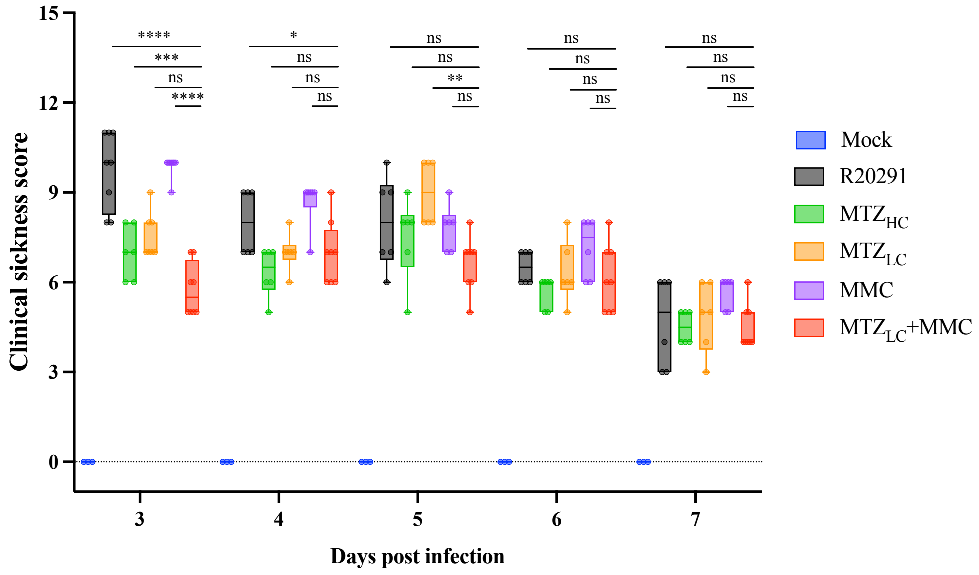

**Supplemental tables**

**Table S1. Bacterial strains used in this study**

| Bacterial strain | Description | Source |
|---|---|---|
| RT027 *C. difficile* strains | | |
| R20291 | $A^+B^+CDT^+$, RT027 | Dr. I-Hsiu Huang |
| CMMC-41 | $A^+B^+CDT^+$, RT027 | Dr. Yuan-Pin Hung |
| CMMC-47 | $A^+B^+CDT^+$, RT027 | Dr. Yuan-Pin Hung |
| NCKUH-93 | $A^+B^+CDT^+$, RT027 | Dr. Yuan-Pin Hung |
| NCKUH-118 | $A^+B^+CDT^+$, RT027 | Dr. Yuan-Pin Hung |
| NTU-50 | $A^+B^+CDT^+$, RT027 | Dr. Yuan-Pin Hung |
| RT078 *C. difficile* clinical strains | | |
| RT078_83 | $A^+B^+CDT^+$, RT078 | Dr. Yuan-Pin Hung |
| RT078_84 | $A^+B^+CDT^+$, RT078 | Dr. Yuan-Pin Hung |
| RT078_85 | $A^+B^+CDT^+$, RT078 | Dr. Yuan-Pin Hung |
| RT078_86 | $A^+B^+CDT^+$, RT078 | Dr. Yuan-Pin Hung |
| RT078_87 | $A^+B^+CDT^+$, RT078 | Dr. Yuan-Pin Hung |
